# Microevolution of acquired colistin resistance in Enterobacteriaceae from ICU patients receiving selective decontamination of the digestive tract

**DOI:** 10.1101/2020.04.26.059691

**Authors:** Axel B. Janssen, Denise van Hout, Marc J.M. Bonten, Rob J.L. Willems, Willem van Schaik

## Abstract

Colistin is an antibiotic that targets the lipopolysaccharides present in the membranes of Gram-negative bacteria. It is used as last-resort drug to treat infections with multidrug-resistant strains. Colistin is also used in selective decontamination of the digestive tract (SDD), a prophylactic therapy used in patients hospitalised in intensive care units (ICUs) to selectively eradicate opportunistic pathogens in the oropharyngeal and gut microbiota. In this study, we aimed to unravel the mechanisms of acquired colistin resistance in Gram-negative opportunistic pathogens obtained from SDD-treated patients.

Routine surveillance of 428 SDD-treated patients resulted in thirteen strains with acquired colistin resistance (*Escherichia coli* n=9; *Klebsiella aerogenes*, n=3; *Enterobacter asburiae*, n=1) from five patients. Genome sequence analysis showed that these isolates represented multiple distinct colistin-resistant clones, but that within the same patients, colistin-resistant strains were clonally related. We identified previously described mechanisms that lead to colistin resistance, i.e. a G53 substitution in the response regulator PmrA/BasR, and the acquisition of the mobile colistin resistance gene *mcr-1*.*1*, but we also observed novel variants of *basR* with an 18-bp deletion, and a G19E substitution in the sensor histidine kinase BasS. We experimentally confirmed these variants to contribute to reduced colistin susceptibility. In a single patient, we observed that colistin resistance in a single *E. coli* clone evolved through two unique variants in *basRS*.

We show that prophylactic use of colistin during SDD can select for colistin resistance in species that are not intrinsically colistin-resistant. This highlights the importance of continued surveillance for the emergence of colistin resistance in patients treated with SDD.

## Introduction

Selective decontamination of the digestive tract (SDD) is a prophylactic antibiotic regimen used in Dutch intensive care units (ICUs) which lowers the mortality in ICU-admitted patients through the selective eradication of opportunistic pathogens in the oropharyngeal and gut microbiota.^1^ One of the targets of SDD are the Enterobacteriaceae, which are collectively responsible for a significant proportion of hospital-acquired infections.^2–6^ In SDD, a combination of the antibiotics colistin and tobramycin and the antifungal amphotericin B, is applied to the digestive tract of ICU patients. In addition, during the first four days of ICU stay, patients are also intravenously administered a biliary-excreted third-generation cephalosporin, contributing to the eradication of Gram-negative pathogens from the gut.^2^ The use of SDD is accompanied by surveillance for potential colonisation of the digestive tract by tobramycin- and/or colistin-resistant Enterobacteriaceae.^7^

Colistin is a cationic cyclic polypeptide with a hydrophobic fatty acid acyl chain that specifically acts on Gram-negative bacteria. Colistin electrostatically interacts with the anionic phosphate groups of the lipid A moiety of lipopolysaccharide (LPS) molecules in the outer leaflet of the outer membrane.^8,9^ Through binding to the phosphate groups, and insertion of its hydrophobic domains, colistin destabilizes the outer membrane. After disruption of the outer membrane, colistin targets the LPS that is resident in the cytoplasmic membrane after its synthesis in the cytoplasm. The destabilization of the cytoplasmic membrane ultimately kills the cell.^10–12^

The most frequent mechanisms of colistin resistance involve the reduction of the anionic charges of lipid A, which reduces the electrostatic interactions between colistin and LPS. This is achieved by the covalent linkage of positively charged groups like phosphoethanolamine or 4-amino-4-deoxy-L-arabinose to the phosphate groups of lipid A.^12^ Addition of these groups to lipid A is achieved by the products of the EptA-, and Arn-operons, respectively.^13^ The transcriptional activity of these operons is controlled by the two-component regulatory systems PhoPQ and PmrAB (BasRS in *Escherichia coli*). Colistin resistance mutations resulting in the permanent activation of these two-component regulatory systems often occur in specific hotspots (e.g. Gly-53 and Arg-81 in BasR/PmrA, and Ala-159 in BasS/PmrB).^14,15^ Surveillance for colistin resistance has recently led to the discovery of novel mechanisms of colistin resistance, including Leu-10 substitutions in BasS/PmrB,^16^ but most importantly the acquisition of *mcr*-genes.^17^ For *E. coli*, the acquisition of *mcr*-carrying mobile genetic elements is a particularly important mechanism through which colistin resistance may occur.^18^ Other mechanisms of acquired colistin resistance in Enterobacteriaceae include the production of capsular polysaccharides,^19^ and efflux pump activation.^20^

The use of SDD is not widely accepted, ^21^ which is at least partially due to the increasing importance of colistin as a last-resort antibiotic for the treatment of infections caused by multidrug-resistant Gram-negative pathogens. ^22,23^ There thus exists an interest in understanding the ability of colistin-resistant strains to emerge and spread during SDD, and to characterise the mechanisms that cause colistin resistance in Enterobacteriaceae from SDD-treated patients. In this work, we analyse thirteen colistin-resistant strains (nine *E. coli* strains, three *Klebsiella aerogenes* strains and one *Enterobacter asburiae* strain) from SDD-treated ICU patients through whole genome sequencing and investigate the mechanisms that have contributed to colistin resistance.

## Material and methods

### Bacterial strains, growth conditions, chemicals, plasmid isolation, and oligonucleotide primers

Colistin-resistant strains were isolated from SDD-treated patients that were hospitalised in the ICU of the University Medical Centre Utrecht, The Netherlands, between July 2018 to January 2019, as previously described.^24^ We also included colistin-susceptible strains of the same species that were isolated from the same patient at the same day from which a colistin-resistant strain was isolated. *E. coli* strain BW25113 and the BW25113-derived Δ*basRS* strain BW27848 from the Keio collection were obtained from the Coli Genetic Stock Center.^25,26^ All strains were grown in Lysogeny Broth (LB; Oxoid, Landsmeer, The Netherlands) at 37°C with agitation at 300 rpm unless otherwise noted. Strains containing pGRG36 were grown at 30°C.^27^ When appropriate, kanamycin (50 mg/L; Sigma-Aldrich, Zwijndrecht, The Netherlands), and ampicillin (100 mg/L; Sigma-Aldrich) were used. Colistin sulphate was obtained from Duchefa Biochemie (Haarlem, The Netherlands). L-(+)-arabinose was obtained from Sigma-Aldrich. Plasmids were purified using the GeneJET Plasmid Miniprep kit (Thermo Fisher Scientific, Landsmeer, The Netherlands). Oligonucleotide primers (Supplemental Table 1) were obtained from Integrated DNA Technologies (Leuven, Belgium).

### Determination of minimal inhibitory concentration

Minimal inhibitory concentrations (MICs) of colistin were determined using a broth microdilution method in line with EUCAST guidelines,^28^ as previously described.^15,29^ The breakpoint values for colistin resistance (MIC ≤ 2 mg/L) in Enterobacteriaceae were obtained from the 2019 European Committee on Antimicrobial Susceptibility Testing guidelines (EUCAST; http://www.eucast.org/clinical_breakpoints/).

### Genomic DNA isolation and whole-genome sequencing

Genomic DNA was isolated, and checked for quality, as described previously.^15^ Sequence libraries for Illumina sequencing were prepared using the Nextera XT kit (Illumina, San Diego, CA) according to the manufacturer’s instructions. Libraries were sequenced on an Illumina NextSeq 500 system with a 300-cycle (2 × 150 bp) NextSeq 500/550 Mid Output v2.5 kit.

### Genome assembly, MLST typing, and identification of antibiotic resistance genes

Illumina sequencing data were assessed (FastQC v0.11.7), and trimmed (nesoni v0.115) for quality, and used for *de novo* genome assembly (SPAdes v3.12.0), as described before.^15,30^ MLST typing was performed using the mlst package v2.10 (https://github.com/tseemann/mlst). Assembled contigs were screened for acquired antibiotic resistance genes using ResFinder 3.2 using standard settings.^31^

### Construction of core genome phylogenetic trees

Genome assemblies of the sequenced strains were aligned with publicly available genomes of the same species obtained from NCBI databases (Supplemental Table 2). Conserved regions of genomes were identified and aligned using ParSNP v1.2.^32^ FigTree v1.4.3 (http://tree.bio.ed.ac.uk/software/figtree/) was used to visualize the phylogenetic tree.

To construct the phylogenetic tree for the clonal strains obtained from patient 31, the genomes of these strains were aligned using snippy v4.4.2 (http://github.com/tseemann/snippy). The alignment was filtered for recombination using gubbins v2.4.1.^33^ Polymorphic sites were subsequently identified through snp-sites v2.5.1.^34^ A phylogenetic tree was constructed using FastTree v2.1.10.^35^ The phylogenetic tree was visualized as described above.

### Determination of SNPs and indels through short-read sequences

Read-mapping of Nesoni-filtered reads was performed using Bowtie2.^36^ SNP and indel-calling was performed using SAMtools 0.1.19 ^37^ through the settings described before.^38^ Identified SNPs and indels were manually linked to features in the Prokka annotation, and inspected for synonymous versus non-synonymous mutations. Identified mutations SNPs and indels were confirmed by performing PCR and Sanger sequencing (Macrogen, Amsterdam, The Netherlands)

### Genome comparisons through Enterobase

To identify mutations that could potentially contribute to colistin resistance in the *E. coli* strains 260 and 263, Raw Illumina sequence reads were uploaded to, and assembled by, Enterobase ^39^ under barcode ESC_OA6301AA and ESC_OA6302AA respectively. The *E. coli* cgMLST scheme was used to define a closest relative in Enterobase and the assemblies of these strains were then used in comparative analyses. Identified sequence variations in the colistin-resistant strains were confirmed by PCR and Sanger sequencing (Macrogen, Amsterdam, The Netherlands)

### Construction of chromosomal *basRS* transgene insertions

Single-copy chromosomal transgene insertion mutants of *basRS*, derived from clinical strains, were constructed in BW27848 using the Tn7-transposon system located on the pGRG36 plasmid,^27,40^ as previously described.^15^

### Determination of maximum specific growth rate

To determine the maximum specific growth rate, a Bioscreen C instrument (Oy Growth Curves AB, Helsinki, Finland) was used. Overnight cultures were used to inoculate 200 µl fresh LB medium 1:1000. Incubation was set at 37°C with continuous shaking set to have maximum amplitude and fastest speed. Growth was observed by measuring the absorbance at 600 nm every 10 minutes.

### Data availability

Sequence data has been deposited in the European Nucleotide Archive (accession number PRJEB34028).

### Statistical analysis

Where applicable, statistical significance was determined using parametric one-way ANOVA tests. Correction for multiple comparison testing was performed for with a Tukey’s (for the all-versus-all comparison of *E. coli* strains of patient 31), or a Dunnett’s (for the *K. aerogenes* strains of patient 37) multiple comparison test. Family-wise significance was defined as a p-value < 0.05.

## Results

### Strains with acquired colistin resistance are rarely isolated during SDD

As described in our previous work, ^24^ 388 Gram-negative strains were isolated from 1105 rectal swabs, from 428 patients receiving SDD. Of these, 102 strains belonged to species that are intrinsically resistant to colistin. The remaining 286 isolates were tested for colistin susceptibility on Sensititre™ FRCOL plates (Thermo Fisher Scientific, Wesel, Germany). A total of ten *E. coli* strains, one *E. asburiae* strain, and three *K. aerogenes* strains were found to be resistant to colistin through this method. We then tested these strains for colistin susceptibility in a standardized broth microdilution assay and all strains were phenotypically resistant to colistin with the exception of *E. coli* strain 89, which was excluded from further analyses (Table 1). Thus, we found thirteen strains (4.5% of the non-intrinsically-resistant isolates) to be colistin-resistant. The colistin-resistant *K. aerogenes* and *E. coli* strains had MIC values up to 32 mg/L colistin. The MIC of the *E. asburiae* strain was found to reach values up to 8192 mg/L colistin. For patients 27 and 37, colistin-susceptible strains of the same species were also isolated from rectal swabs during surveillance (Figure 1).

**Table 1.** Strains isolated in this study and relevant metadata. The species of each isolate was determined by MALDI-TOF on a Bruker microflex system (Leiderdorp, The Netherlands). The MIC value of the broth microdilution method represents the median value of three independent replicates of colistin susceptibility testing, performed in triplicate.

| Strain | Date of isolation | Patient | Species | Colistin MIC (mg/L) |
| --- | --- | --- | --- | --- |
| 24 | 6 August 2018 | 37 | *Klebsiella aerogenes* | 0.25 |
| 25 | 6 August 2018 | 37 | *Klebsiella aerogenes* | 32 |
| 26 | 9 August 2018 | 37 | *Klebsiella aerogenes* | 32 |
| 27 | 9 August 2018 | 37 | *Klebsiella aerogenes* | 32 |
| 32 | 7 September 2018 | 307 | *Enterobacter asburiae* | 8192 |
| 89 | 26 July 2018 | 337 | *Escherichia coli* | 0.25 |
| 137 | 30 August 2018 | 27 | *Escherichia coli* | 0.125 |
| 138 | 30 August 2018 | 27 | *Escherichia coli* | 8 |
| 260 | 1 November 2018 | 31 | *Escherichia coli* | 16 |
| 262 | 1 November 2018 | 31 | *Escherichia coli* | 16 |
| 263 | 1 November 2018 | 311 | *Escherichia coli* | 4 |
| 274 | 5 November 2018 | 31 | *Escherichia coli* | 16 |
| 281 | 8 November 2018 | 31 | *Escherichia coli* | 16 |
| 292 | 14 November 2018 | 31 | *Escherichia coli* | 16 |
| 296 | 14 November 2018 | 31 | *Escherichia coli* | 16 |
| 297 | 14 November 2018 | 31 | *Escherichia coli* | 16 |

**Figure 1.**
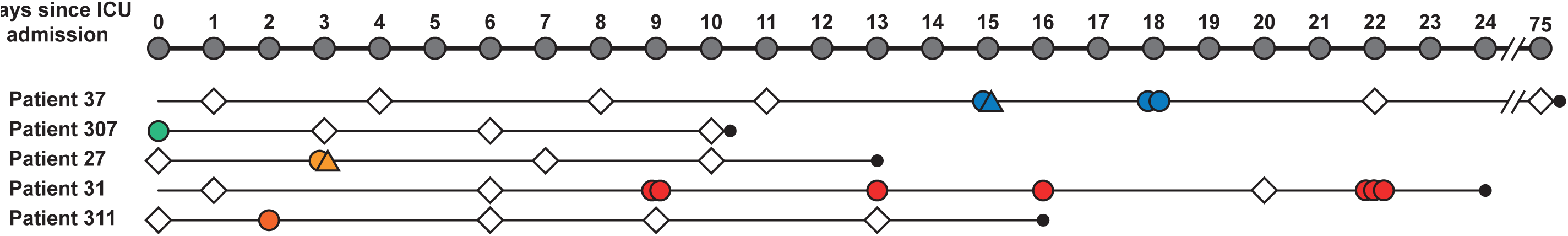
Timeline of rectal swabs collected from SDD-treated ICU patients and isolation of colistin-susceptible, or colistin-resistant strains. Filled circle: isolation of colistin-resistant strain from rectal swab. Filled triangle; a colistin-susceptible strain of the same species as the colistin-resistant strain was isolated from the rectal swab. Open diamonds, no naturally colistin-susceptible Gram-negative bacteria were isolated. The symbols are colour coded according to the species of the isolated strain: *Klebsiella aerogenes*, blue; *Enterobacter asburiae*, green; *Escherichia coli*, yellow, orange and red. Multiple symbols on the same day indicate the isolation of multiple strains from the same swab. The length of ICU admission is indicated by a line, and discharge is indicated by a circle at the end of the line.

In total, five of the 428 patients (1.2%) tested positive for an isolate with acquired colistin resistance. Of the ten *E. coli* strains, seven were isolated from one patient, the remaining three strains originated from three other patients. The three *K. aerogenes* strains were isolated from a single patient. None of the patients carried multiple colistin-resistant species. Of the five patients from whom a strain with acquired colistin resistance was isolated, only patient 307 carried a colistin-resistant strain (*E. asburiae*) at the start of ICU admission, suggesting that this patient had acquired the colistin-resistant strain prior to SDD treatment (Figure 1). This strain was no longer present on any of the following time-points. Patients 311 and 27 were transiently colonised by colistin-resistant strains at 2 and 3 days after ICU admission, respectively. In patient 37 colistin-resistant *K. aerogenes* were detected at day 15, and day 18 of ICU hospitalisation, while strains with acquired colistin resistance were not cultured during subsequent screening from day 22 to discharge from the ICU on day 75. Colistin-resistant *E. coli* were first detected in patient 31 at day 9 of ICU hospitalisation and remained colonised with colistin-resistant *E. coli*, until the patient was lost to follow up at day 24, with the exception of a negative culture result on day 20.

### Colistin-resistant strains from ICU patients have a diverse genetic background and carry a variety of acquired antibiotic-resistance genes

Clonal relatedness of colistin-resistant *E. coli* and *K. aerogenes* strains colonising the ICU patients was assessed by constructing core genome phylogenies, using the genome sequences generated in this study and a collection of publicly available genomes. We observed that three distinct clones of colistin-resistant *E. coli* colonised three individual patients (Figure 2A). The colistin-susceptible *E. coli* strain 137 from patient 27 did not cluster with colistin-resistant *E. coli* strain 138 isolated from the same patient. All *K. aerogenes* strains isolated from patient 37 belonged to a single clone (Figure 2B).

**Figure 2.**
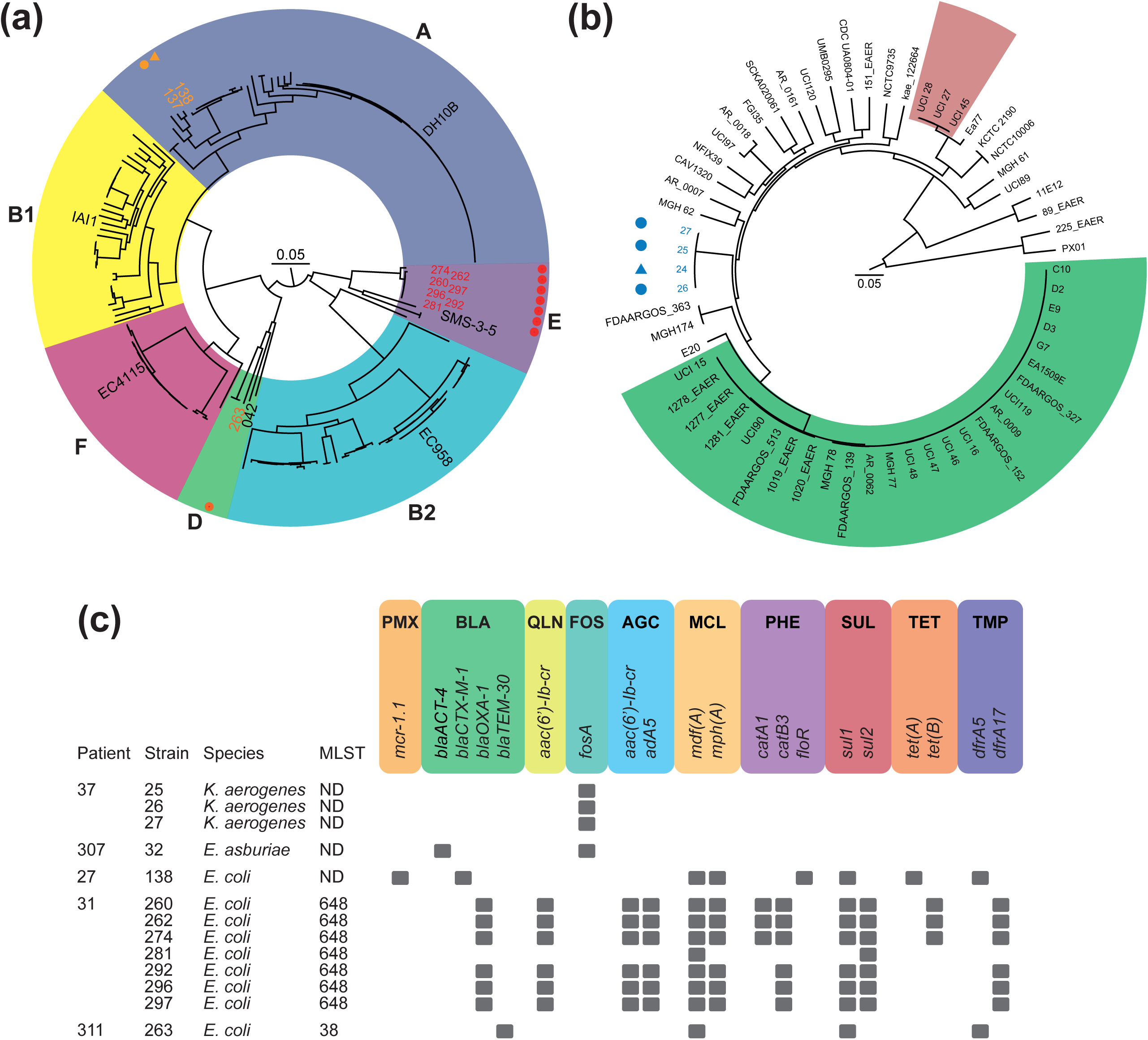
Phylogeny and acquired resistance genes of colistin-resistant *E. coli* and *K. aerogenes*. A) The phylogenetic tree of *E. coli* represents the core genome alignment (2.1 Mbp) of 198 genomes (178 genomes from public databases, the genomes of the nine colistin-resistant strains, and the two colistin-susceptible strains). One representative reference strain per *E. coli* phylogroup is indicated ^69^. Phylogroup A is coloured dark blue; B1, yellow; B2, light blue; D, green; E, purple; F, pink. Colistin-susceptible and -resistant strains are indicated by filled triangles and circles respectively. The studied strains are highlighted with colours corresponding to Figure 1. B) The phylogenetic tree of *K. aerogenes* represents a core genome alignment (1.6 Mbp) of 56 publicly available genomes, and the three colistin-resistant strains described in this study. Colistin-susceptible and -resistant strains are indicated by filled triangles and circles respectively. The studied strains are highlighted with colours corresponding to Figure 1. The genomes of the dominant *K. aerogenes* ST4 and ST93 lineages associated with infections have an red and green background respectively ^70^. C) Acquired antibiotic resistance genes of colistin-resistant strains. Strains are grouped according to the patient from which they were isolated. Species and MLST type are indicated per strain. Antibiotic resistance genes in the genomes of the colistin-resistant strains were detected by ResFinder 3.2 ^31^. Classes of antibiotic resistance genes are abbreviated as follow: PMX, polymyxin resistance; BLA, beta-lactam resistance; QLN, quinolone resistance; FOS, fosfomycin resistance; AGC, aminoglycoside resistance; MCL, macrolide, lincosamide, and streptogramin B resistance; PHE, phenicol resistance; SUL, sulfonamide resistance; TET, tetracycline resistance; TMP, trimethoprim resistance. N.D.; not determined.

We screened the assembled genomes of the colistin-resistant strains for acquired antibiotic resistance genes (Figure 2C). We identified that strain *E. coli* strain 138 was positive for the *mcr-1*.*1* gene, which was the sole antibiotic resistance gene on a 59.5 kbp contig that contained an IncI2-type replicon. The sequence of this contig shared 99% identity with plasmid sequences obtained from multiple sources, including from a *Salmonella enterica* subsp. e*nterica* serovar Typhimurium strain from a patient from China (accession number MH522416.1),^41^ and an *E. coli* strain isolated from chicken faeces in Thailand (accession number MG557851.1),^42^ illustrating the global spread of this plasmid. In addition, we found that the colistin-resistant *E. coli* strains carried between 2 and 12 antibiotic resistance genes, including genes conferring resistance to aminoglycosides and β-lactams. The *K. aerogenes* strains only carried the fosfomycin resistance gene *fosA*, and the single *E. asburiae* strain had *fosA* and the AmpC-type β-lactamase *blaACT-4*.

### In-patient microevolution of colistin resistance in *Escherichia coli* during SDD

To investigate the microevolution towards colistin resistance during gut colonisation, we investigated the genetic diversity between the four clonally related *K. aerogenes* strains from patient 37, and the seven colistin-resistant *E. coli* strains isolated from patient 31. By comparing the genome of the single colistin-susceptible *K. aerogenes* strain with the genomes of the colistin-resistant strains, we determined that all colistin-resistant strains had a G53S substitution in the PmrA transcriptional regulator of the PmrAB two-component regulatory system. This substitution has previously been described to cause colistin resistance in *K. aerogenes*.^43^ In addition, we observed an insertion of a single guanine nucleotide in the gene encoding the sulphate adenylyltransferase subunit 2 CysD, which is involved in sulphate assimilation. CysD has not been previously described in relation to colistin resistance. No mutations differentiating the three colistin-resistant strains were observed.

The seven clonally related colistin-resistant *E. coli* isolates from patient 31 were obtained on four separate days, over a two-week period (Table 1, Figure 2A). Within these strains, we observed that the strains isolated on 1 and 5 November 2018 (strains 260, 262, and 274) had a G53A substitution in the BasR transcriptional regulator of the BasRS two-component regulatory system. This substitution has previously been experimentally proven to contribute to colistin resistance in *E. coli*.^15^ The strains isolated after November 5, 2018 (strains 281, 292, 296, and 297) however, did not encode the G53A BasR substitution. Instead, we observed an 18-bp deletion (nucleotides 10 through 27) in *basS*, leading to the deletion of six amino acids (4 through 9) at the N-terminal end of BasS.

Through phylogenetic analysis of these seven clonally related strains, we observed that the strains carrying the G53A BasR substitution, and those with the 18-bp deletion in *basS*, were intermingled (Figure 3A). Three strains that were all isolated on 14 November 2018 (strains 292, 296, and 297) had a mucoid phenotype (Figure 3B) and carried an I424S substitution in YrfF that was absent in the non-mucoid strains. YrfF is a homolog of IgaA in *Salmonella*, and functions as a negative regulator of the Rcs phosphorelay system, and is thus involved in regulation of capsule production.^44^

**Figure 3.**
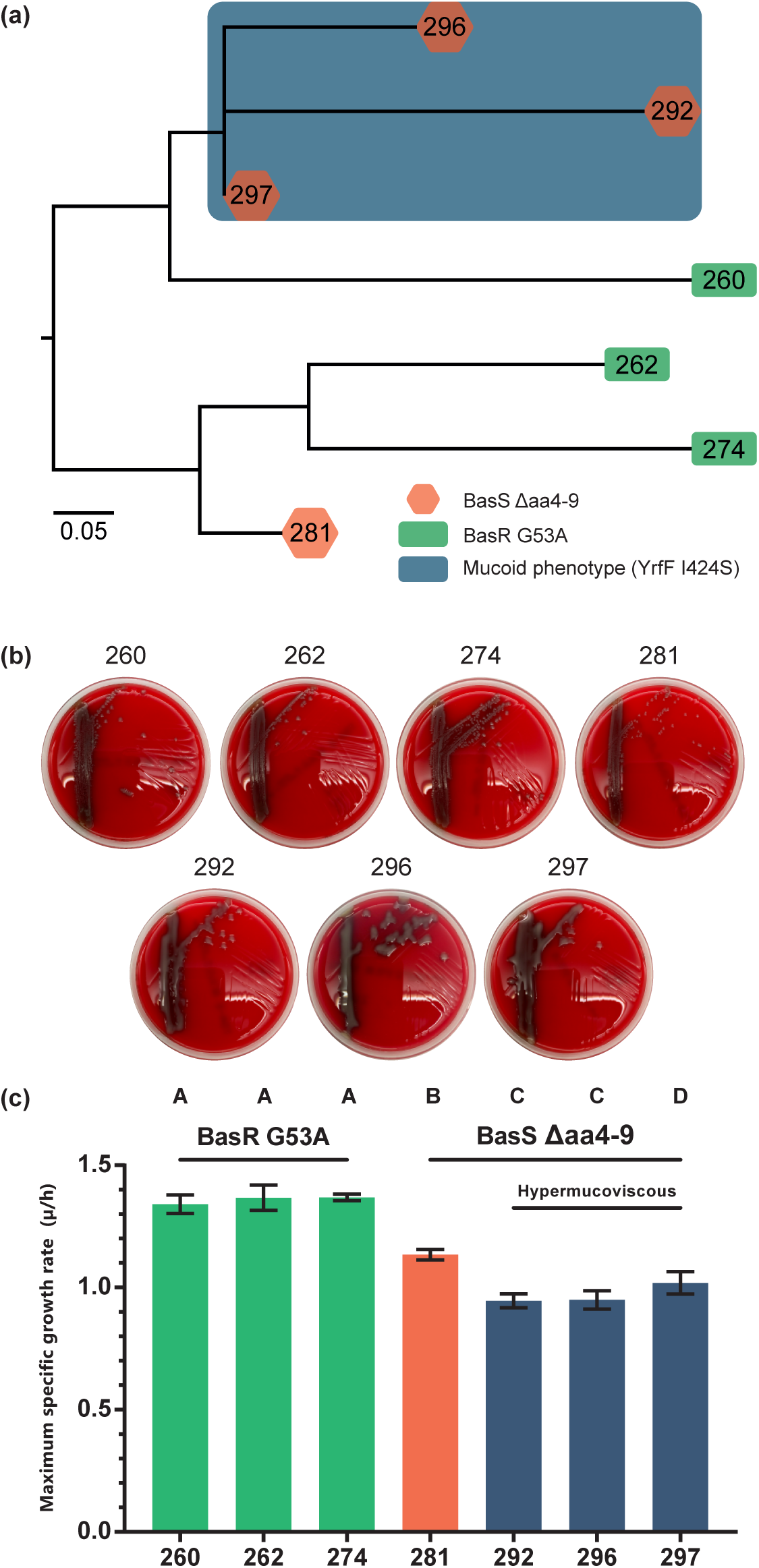
*In vivo* microevolution of colistin-resistant *E. coli* strains isolated from patient 31. A) Mid-point rooted phylogenetic tree representing a recombination-filtered core genome alignment of the strains isolated from patient 31. Branches with the BasR G53A substitution, the BasS Δaa4-9 deletion, and the I424S substitution in YrfF are indicated through a green, orange, and purple background respectively. Other mutations are indicated next to the relevant branch. B) Growth phenotypes of the *E. coli* strains isolated from patient 31 on TSA plates with 5% sheep blood after overnight growth at 37°C. C) Maximum specific growth rate of colistin-resistant *E. coli* strains isolated from patient 31. The values presented represent mean with standard deviation, of three independent experiments, performed in duplicate. Statistical significance testing was performed using a parametric one-way ANOVA test with Tukey’s multiple comparisons test. Family-wise significance was defined as a p-value < 0.05. The growth rates of strains marked by the same letter differ statistically significantly from those with other letters.

To establish the impact of colistin resistance on fitness, we determined the strains’ maximum specific growth rate. Among the strains from patient 31, we observed a reduction of about 15% in maximum specific growth rate in the strains harbouring the 18-bp deletion in *basS*, compared to those harbouring the G53A BasR substitution (Figure 3C). A similar decrease in maximum specific growth rate is observed in the strains that have the I424S substitution in YrfF. For the *K. aerogenes* strains isolated from patient 37 that had the G53S PmrA substitution, we did not observe a change in the maximum specific growth rate (Supplemental Figure 1).

### Mutations in *basS* contribute to reduced susceptibility to colistin

To identify mutations that potentially contributed to colistin resistance in *E. coli* strain 263, for which we lacked an isogenic, colistin-susceptible counterpart, we used the Enterobase database to identify the strain (eo2071, which was not reported to be colistin-resistant; Enterobase barcode ESC_BA7113AA) that is most closely related to strain 263. We then compared the sequence of *basRS* of strain eo2071 to *basRS* of strain 263, leading to the identification of a G19E substitution in BasS in the latter strain.

We next aimed to investigate the relevance of the 18-bp deletion in *basS* and the mutation leading to the G19E substitution for colistin resistance in *E. coli*. Due to the multidrug-resistant nature of the colistin-resistant strains, we constructed chromosomal integration mutants of the genes encoding the mutated BasRS two-component regulatory system in the *att*Tn7 site in the BW25113 derived Δ*basRS* strain BW27848, as described previously. ^15^ MIC determinations of the chromosomal integration mutants showed that introduction of the *basRS* alleles of the colistin-resistant strains led to reduced susceptibility to colistin (Table 2). The MIC value of the BW27848 strain with the insertion of *basRS* encoding the 18-bp deletion in *basS* increased 4-fold. The MIC value of the BW27848 strain with the *basRS* encoding the G19E substitution in *basR* increased 2-fold. Restoring the deleted 18 base-pairs to *basS*, or reversing the G19E BasR substitution, returned colistin susceptibility to BW25113 levels.

**Table 2.** Colistin MICs of strains generated in this study. *E. coli* strain BW27848 is the Δ*basRS* mutant of BW25113.^26^ The *basRS* alleles of colistin-resistant strains from this study were inserted into the *att*Tn7 site of BW27848. The addition of “m” to a strain name indicates that the construct has been modified through inverse PCR site-directed mutagenesis to reverse the mutation associated with colistin resistance.

| Strain | Colistin MIC (mg/L) |
| --- | --- |
| BW25113 | 0.125 |
| BW27848 | 0.25 |
| BW25113::Tn7 Empty | 0.125 |
| BW27848::Tn7 Empty | 0.125 |
| BW27848::Tn7 BW25113 | 0.125 |
| BW27848::Tn7 281 | 1 |
| BW27848::Tn7 281m | 0.125 |
| BW27848::Tn7 263 | 0.25 |
| BW27848::Tn7 263m | 0.125 |

## Discussion

In this study, we genomically characterized strains with acquired colistin resistance from ICU patients treated with SDD. Thirteen strains, from three species with acquired colistin resistance were isolated. The carriage rate of strains with acquired colistin resistance among SDD-treated ICU patients (1.2%) was similar to rates previously reported for Dutch ICUs. ^6,15,45^ We found multiple distinct clones of colistin-resistant strains among patients. Colistin resistance in the *E. coli* strains could be explained by the acquisition of *mcr-1*.*1*, a G53A or G19E substitution in BasR, or an 18-bp deletion in *basS*. The 18-bp deletion in *basS* was associated with a loss of fitness. Colistin resistance in the *K. aerogenes* strains was associated with a G53S substitution in PmrA. The mechanism of colistin resistance in the *E. asburiae* strain was not identified.

Through longitudinal sampling, we are able to observe selection and microevolutionary processes related to prolonged exposure to colistin. We found that colistin resistance in a clonal *E. coli* population may emerge through different mutational trajectories in *basRS*, as observed in the strains obtained from patient 31. The placement of strains with different mutations in *basRS* (either causing a G53A BasR substitution or a 18-bp deletion in *basS*) on distinct branches of the phylogenetic tree, suggests that strains with both mutations have emerged independently. The observation of two distinct variants in *basRS* that contribute to colistin resistance in a clonal *E. coli* population, reflects the ability of *E. coli* to quickly adapt to novel niches and selective pressures through multiple evolutionary pathways.^46–48^ This *E. coli* clone was likely able to persistently colonise the patient because it carried mechanisms that confer resistance to all antibiotics (cephalosporins, tobramycin, and colistin) used in SDD. Through longitudinal sampling of isogenic colistin-susceptible and colistin-resistant *E. coli* strains from patients 27 and 37, we observed that colistin resistance emerged *de novo* in these strains. For the strains isolated from patients 307 and 311, we cannot exclude the possibility that these strains could have already acquired colistin resistance before they colonised the patient.

We observed transient colonisation by an *mcr-1*.*1* carrying *E. coli* in a single patient receiving SDD. Transient gut colonisation by *mcr-*carrying *E. coli* has previously been observed in the context of international travel, and may reflect the reduced fitness of *E. coli* strains carrying *mcr*-genes.^49–51^ High-level colistin resistance was observed in an *Enterobacter asburiae* strain, a member of the *E. cloacae* complex,^52^ and a clonal population of colistin-resistant *K. aerogenes*. A limited number of high-level resistant *E. asburiae*,^53–57^ and *K. aerogenes* ^43,58–60^ strains have been described before. However, the exact mechanisms through which colistin resistance evolves in these species remain poorly studied.^43,61^

Prolonged exposure to antibiotics will select for resistance, which generally comes at a fitness cost. Fitness of an antibiotic-resistant clone can subsequently increase due to the accumulation of compensatory mutations.^62^ Interestingly, we observed a reduction of the maximum specific growth rate in the *E. coli* strains in which *yrfF* was mutated, which likely contributed to the mucoid phenotype in these strains. The increased biosynthesis of capsular polysaccharides is likely to come at a cost that will negatively impact maximum specific growth rate. While mucoidy has been linked to colistin resistance in *K. pneumoniae*, and *Neisseria meningitides*,^19,63,64^ we did not observe a difference in the colistin MICs of clonally related non-mucoid and mucoid *E. coli* strains. Mucoidy in *E. coli* is a relatively poorly understood phenotype, but it is likely to contribute to increased survival upon humoral and cellular immune responses. ^65,66^

In this study, we find that Gram-negative opportunistic pathogens carried in the gut of patients can acquire colistin resistance, either through mutation of genes that regulate lipid A modifications or by the acquisition of the *mcr-1*.*1* gene. However, the low prevalence of colistin-resistant strains in ICU patients suggests that the evolution of colistin resistance is currently of minor concern for the implementation of SDD in Dutch hospitals. As the prevalence of multidrug-resistant Gram-negative bacteria in the Netherlands is low, strains colonising patients will be generally susceptible to one or more of the antibiotics used in SDD. Indeed, in four patients, strains with acquired colistin resistance were rapidly eradicated from the gut. However, the long-term colonisation of patient 31 with an *E. coli* clone that is colistin resistant and carries genes conferring resistance to the other antibiotics used in SDD, indicates that SDD can select for multidrug-resistant Gram-negative bacteria in the gut of ICU patients. The risk of the emergence of colistin resistance in the patient gut may be more pronounced in countries where higher rates of circulating antibiotic resistant bacteria are observed, or in settings with failing infection control.^6,22,67,68^ Continuous surveillance is thus vital to thwart selection and spread of multidrug-resistant strains upon SDD. Further studies are required to better understand the diversity of mechanisms of acquired colistin resistance in clinical Enterobacteriaceae isolates, particularly in species like *E. asburiae*, and *K. aerogenes* in which colistin resistance mechanisms have so far been poorly studied.

## Acknowledgements

We thank Johanna C. Braat, Ellen C. Brouwer, Malbert R.C. Rogers, Moniek Salomons, and the Department of Genetics at the Wilhelmina Children’s Hospital, for their expertise on Illumina NextSeq sequencing. We thank Rob Rentenaar, and Judith Vlooswijk for providing the strains that were studied here.

## Transparency

The authors disclose no conflicts of interest.

## Author contributions

A.B.J. conceived and designed experiments, performed experiments, analysed data, and wrote the manuscript. D.v.H. reviewed patients records and microbiological data. M.J.M.B. wrote the manuscript. R.J.L.W. wrote the manuscript. W.v.S. conceived and designed experiments, wrote the manuscript, and supervised the study. All authors reviewed and approved the final version of the manuscript.

## Supplemental materials

**Supplemental Figure 1. Maximum specific growth rate of *K. aerogenes* strains**. Maximum specific growth rate of the colistin-susceptible, and colistin-resistant *K. aerogenes* strains isolated from patient 37. The values presented represent the means with standard deviations of three independent experiments performed in triplicate. Statistical significance testing was performed by comparing the maximum specific growth rate of the colistin-susceptible strain 24, with those of the colistin-resistant strains through a parametric one-way ANOVA test, with a Dunnett’s multiple comparisons test. Family-wise significance was defined as a p-value < 0.05. No statistical significant differences were observed between the maximum specific growth rates of the strains.

**Supplemental Table 1. Oligonucleotides used in this study**.

**Supplemental Table 2. Strains used for phylogenetic analyses**. A total of 178 *E. coli* and 56 *K. aerogenes* genome sequences obtained from NCBI databases were used for construction of phylogenetic trees. When multiple assemblies had the same strain name, a numerical indicator was added.

