## Supplemental Table 1 for "Microevolution of acquired colistin resistance in Enterobacteriaceae from ICU patients receiving selective decontamination of the digestive tract"

| **Name** | **Sequence (5'- 3')^a^** | **Reference** |
| --- | --- | --- |
| M13 Fwd | TGT AAA ACG ACG GCC AGT | This study |
| M13 Rev | CAG GAA ACA GCT ATG ACC | This study |
| BW25113 glmS Up | ATA TTC AGT CAA TTA CAA ACA TTA | ^15^ |
| BW25113 glmS Down | CGA TCT TCT ACA CCG TTC | ^15^ |
| BasRS sequencing Fwd | AAA GCC CGT ATC CGC AC | ^15^ |
| BasRS sequencing Rev | GAT CTC ACG CAT GAT GTG GC | ^15^ |
| Tn7R inward | GAA ATC AGT CCA GTT ATG CTG TG | ^15^ |
| Tn7L inward | GGG TGT AGC GTC GTA AGC TAA T | ^15^ |
| BasRS genes Fwd | ATG AAA ATT CTG ATT GTT GAA GA | ^15^ |
| BasRS genes Rev | GTT TAG CGT GCT GGT | This study |
| BasRS promoter Fwd | CTT CCT CTA CTG CAT CTG GG | ^15^ |
| BasRS promoter Rev | CAC GGT GTT TCC ATC GA | ^15^ |
| BasRS genes overlap Fwd | TGC GCA CTT TGT TCG ATG GAA ACA CCG TGA TGA AAA TTC TGA TTG TTG | This study |
| BasRS promotor Fwd NotI | TAT CCT GCG GCC GCC TTC CTC TAC TGC ATC TGG G | ^15^ |
| BasRS genes Rev XhoI | TAT CCC CTC GAG GTT TAG CGT GCT GGT GGT | This study |
| BasRS promotor Fwd NotI short | TAT CCT GCG GCC GCC | ^15^ |
| BasRS genes Rev XhoI short | TAT CCC CTC GAG GTT TA | This study |
| BasS Δ18bp restore Fwd | AAA ACG CAT CAG ATT CAA TTA GTT TTC CTC ATT | This study |
| BasS Δ18bp restore Rev | CTG CGC CGA CCA ATA TCG CTG CGC CAA CGG CTG ATA TT | This study |
| BasR E19G F | ACG GCT GAT ATT GAC CAT TGG GGC CAT TT TGT TG | This study |
| BasR E19G R | TGG CGC AGC GAT ATT GGT CGG CG | This study |
| Tn7L | ATT AGC TTA CGA CGC TAC ACC C | ^40^ |
| pGRG36 check with Tn7L | ATA TGC ACA GAT GAA AAC GGT G | ^15^ |

^a^ Restriction sites are underlined.
