## Supplemental Table 2 for "Microevolution of acquired colistin resistance in Enterobacteriaceae from ICU patients receiving selective decontamination of the digestive tract"

| **Strain** | **Genbank accession number** | **Species** |
| --- | --- | --- |
| 42 | GCA_000027125.1 | *Escherichia coli* |
| 536 | GCA_000013305.1 | *Escherichia coli* |
| 789 | GCA_000819645.1 | *Escherichia coli* |
| 1303 | GCA_000829985.1 | *Escherichia coli* |
| 6409 | GCA_000814145.2 | *Escherichia coli* |
| 11128 | GCA_000010765.1 | *Escherichia coli* |
| 11368 | GCA_000091005.1 | *Escherichia coli* |
| 12009 | GCA_000010745.1 | *Escherichia coli* |
| 55989 | GCA_000026245.1 | *Escherichia coli* |
| 180-PT54 | GCA_001650275.1 | *Escherichia coli* |
| 2009C-3133 | GCA_001420955.1 | *Escherichia coli* |
| 2009EL-2050 | GCA_000299255.1 | *Escherichia coli* |
| 2009EL-2071 | GCA_000299475.1 | *Escherichia coli* |
| 2011C-3493 | GCA_000299455.1 | *Escherichia coli* |
| 2011C-3911 | GCA_001644725.1 | *Escherichia coli* |
| 2012C-4227 | GCA_001420935.1 | *Escherichia coli* |
| 2013C-4465 | GCA_001644745.1 | *Escherichia coli* |
| 28RC1 | GCA_001612475.1 | *Escherichia coli* |
| 644-PT8 | GCA_001650295.1 | *Escherichia coli* |
| 94-3024 | GCA_000801185.2 | *Escherichia coli* |
| ABU 83972 | GCA_000148365.1 | *Escherichia coli* |
| ACN001 | GCA_001051135.1 | *Escherichia coli* |
| ACN002 | GCA_001515725.1 | *Escherichia coli* |
| APEC IMT5155 | GCA_000813165.1 | *Escherichia coli* |
| APEC O1 | GCA_000014845.1 | *Escherichia coli* |
| APEC O78 | GCA_000332755.1 | *Escherichia coli* |
| ATCC 25922 | GCA_000743255.1 | *Escherichia coli* |
| ATCC 8739 | GCA_000019385.1 | *Escherichia coli* |
| B7A | GCA_000725265.1 | *Escherichia coli* |
| BL21 (TaKaRa) | GCA_000833145.1 | *Escherichia coli* |
| BL21(DE3) | GCA_000009565.2 | *Escherichia coli* |
| BL21(DE3) | GCA_000022665.2 | *Escherichia coli* |
| BL21-Gold(DE3)pLysS AG | GCA_000023665.1 | *Escherichia coli* |
| C227-11 | GCA_000986765.1 | *Escherichia coli* |
| C2566 | GCA_001559615.1 | *Escherichia coli* |
| C3026 | GCA_001559675.1 | *Escherichia coli* |
| C3029 | GCA_001559635.1 | *Escherichia coli* |
| C321.deltaA | GCA_000474035.1 | *Escherichia coli* |
| C41(DE3) | GCA_000830035.1 | *Escherichia coli* |
| C43(DE3) | GCA_001039415.1 | *Escherichia coli* |
| CB9615 | GCA_000025165.1 | *Escherichia coli* |
| CD306 | GCA_001513615.1 | *Escherichia coli* |
| CE10 | GCA_000227625.1 | *Escherichia coli* |
| CFSAN029787 | GCA_001007915.1 | *Escherichia coli* |
| CFT073 | GCA_000007445.1 | *Escherichia coli* |
| CI5 | GCA_000971615.1 | *Escherichia coli* |
| clone D i14 | GCA_000233895.1 | *Escherichia coli* |
| clone D i2 | GCA_000233875.1 | *Escherichia coli* |
| CQSW20 | GCA_001455385.1 | *Escherichia coli* |
| DH1 #1 | GCA_000023365.1 | *Escherichia coli* |
| DH1 #2 | GCA_000270105.1 | *Escherichia coli* |
| DH1Ec095 | GCA_001183645.1 | *Escherichia coli* |
| DH1Ec104 | GCA_001183665.1 | *Escherichia coli* |
| DH1Ec169 | GCA_001183685.1 | *Escherichia coli* |
| DHB4 | GCA_001559655.1 | *Escherichia coli* |
| E2348/69 | GCA_000026545.1 | *Escherichia coli* |
| E24377A | GCA_000017745.1 | *Escherichia coli* |
| EC4115 | GCA_000021125.1 | *Escherichia coli* |
| EC958 | GCA_000285655.3 | *Escherichia coli* |
| ECC-1470 | GCA_000831565.1 | *Escherichia coli* |
| Eco889 | GCA_001663475.1 | *Escherichia coli* |
| Ecol_448 | GCA_001618365.1 | *Escherichia coli* |
| Ecol_732 | GCA_001617565.1 | *Escherichia coli* |
| Ecol_743 | GCA_001618325.1 | *Escherichia coli* |
| Ecol_745 | GCA_001618345.1 | *Escherichia coli* |
| ECONIH1 | GCA_000784925.1 | *Escherichia coli* |
| EDL933 | GCA_000732965.1 | *Escherichia coli* |
| ER1821R | GCA_001663075.1 | *Escherichia coli* |
| ER2796 | GCA_000800215.1 | *Escherichia coli* |
| ER3413 | GCA_000800765.1 | *Escherichia coli* |
| ER3435 | GCA_000974885.1 | *Escherichia coli* |
| ER3440 | GCA_000974465.1 | *Escherichia coli* |
| ER3445 | GCA_000974535.1 | *Escherichia coli* |
| ER3446 | GCA_000974825.1 | *Escherichia coli* |
| ER3454 | GCA_000974405.1 | *Escherichia coli* |
| ER3466 | GCA_000974575.1 | *Escherichia coli* |
| ER3475 | GCA_000974865.1 | *Escherichia coli* |
| ER3476 | GCA_000974505.1 | *Escherichia coli* |
| ETEC H10407 | GCA_000210475.1 | *Escherichia coli* |
| FRIK2069 | GCA_001651925.1 | *Escherichia coli* |
| FRIK2455 | GCA_001651965.1 | *Escherichia coli* |
| FRIK2533 | GCA_001651945.1 | *Escherichia coli* |
| G749 | GCA_001566635.1 | *Escherichia coli* |
| HS | GCA_000017765.1 | *Escherichia coli* |
| HUSEC2011 | GCA_000967155.1 | *Escherichia coli* |
| IAI1 | GCA_000026265.1 | *Escherichia coli* |
| IAI39 | GCA_000026345.1 | *Escherichia coli* |
| IHE3034 | GCA_000025745.1 | *Escherichia coli* |
| JEONG-1266 | GCA_001558995.2 | *Escherichia coli* |
| JJ1886 | GCA_000493755.1 | *Escherichia coli* |
| JJ1887 | GCA_001593565.1 | *Escherichia coli* |
| JJ1897 | GCA_001513655.1 | *Escherichia coli* |
| JJ2434 | GCA_001513635.1 | *Escherichia coli* |
| JW5437-1 substr. MG1655 | GCA_001566335.1 | *Escherichia coli* |
| K-12 substr. AG100 | GCA_000981485.1 | *Escherichia coli* |
| K-12 substr. BW25113 | GCA_000750555.1 | *Escherichia coli* |
| K-12 substr. BW2952 | GCA_000022345.1 | *Escherichia coli* |
| K-12 substr. DH10B | GCA_000019425.1 | *Escherichia coli* |
| K-12 substr. GM4792 #1 | GCA_001020945.2 | *Escherichia coli* |
| K-12 substr. GM4792 #2 | GCA_001021005.2 | *Escherichia coli* |
| K-12 substr. HMS174 | GCA_000953515.1 | *Escherichia coli* |
| K-12 substr. MC4100 | GCA_000499485.1 | *Escherichia coli* |
| K-12 substr. MDS42 | GCA_000350185.1 | *Escherichia coli* |
| K-12 substr. MG1655 #1 | GCA_000005845.2 | *Escherichia coli* |
| K-12 substr. MG1655 #2 | GCA_000801205.1 | *Escherichia coli* |
| K-12 substr. MG1655 #3 | GCA_001308065.1 | *Escherichia coli* |
| K-12 substr. MG1655 #4 | GCA_001544635.1 | *Escherichia coli* |
| K-12 substr. MG1655_TMP32XR1 | GCA_001308125.1 | *Escherichia coli* |
| K-12 substr. MG1655_TMP32XR2 | GCA_001308165.1 | *Escherichia coli* |
| K-12 substr. RV308 | GCA_000952955.1 | *Escherichia coli* |
| K-12 substr. W3110 | GCA_000010245.1 | *Escherichia coli* |
| KLY | GCA_000725305.1 | *Escherichia coli* |
| KO11 | GCA_000147855.3 | *Escherichia coli* |
| KO11FL | GCA_000258025.1 | *Escherichia coli* |
| LF82 | GCA_000284495.1 | *Escherichia coli* |
| LY180 | GCA_000468515.1 | *Escherichia coli* |
| MNCRE44 | GCA_000931565.1 | *Escherichia coli* |
| MRE600 | GCA_001542675.2 | *Escherichia coli* |
| MVAST0167 | GCA_001566655.1 | *Escherichia coli* |
| NA114 | GCA_000214765.2 | *Escherichia coli* |
| NCM3722 | GCA_001043215.1 | *Escherichia coli* |
| NGF1 | GCA_001660585.1 | *Escherichia coli* |
| Nissle 1917 | GCA_000714595.1 | *Escherichia coli* |
| NRG 857C | GCA_000183345.1 | *Escherichia coli* |
| P12b | GCA_000257275.1 | *Escherichia coli* |
| PCN033 | GCA_000219515.3 | *Escherichia coli* |
| PCN061 | GCA_001029125.1 | *Escherichia coli* |
| REL606 | GCA_000017985.1 | *Escherichia coli* |
| RM12579 | GCA_000245515.1 | *Escherichia coli* |
| RM12581 | GCA_000671295.1 | *Escherichia coli* |
| RM12761 | GCA_000662395.1 | *Escherichia coli* |
| RM13514 | GCA_000520035.1 | *Escherichia coli* |
| RM13516 | GCA_000520055.1 | *Escherichia coli* |
| RM9387 | GCA_000801165.1 | *Escherichia coli* |
| RR1 | GCA_001276585.1 | *Escherichia coli* |
| RS218 | GCA_000800845.2 | *Escherichia coli* |
| S51 | GCA_001660565.1 | *Escherichia coli* |
| S88 | GCA_000026285.1 | *Escherichia coli* |
| Sakai substr. RIMD 0509952 | GCA_000008865.1 | *Escherichia coli* |
| Sanji | GCA_001610755.1 | *Escherichia coli* |
| Santai | GCA_000827105.1 | *Escherichia coli* |
| SaT040 | GCA_001566615.1 | *Escherichia coli* |
| SE11 | GCA_000010385.1 | *Escherichia coli* |
| SE15 | GCA_000010485.1 | *Escherichia coli* |
| SEC470 | GCA_000987875.1 | *Escherichia coli* |
| SF-088 | GCA_001280325.1 | *Escherichia coli* |
| SF-166 | GCA_001280385.1 | *Escherichia coli* |
| SF-173 | GCA_001280405.1 | *Escherichia coli* |
| SF-468 | GCA_001280345.1 | *Escherichia coli* |
| SMS-3-5 | GCA_000019645.1 | *Escherichia coli* |
| SQ110 | GCA_000988425.1 | *Escherichia coli* |
| SQ171 | GCA_000988445.1 | *Escherichia coli* |
| SQ2203 | GCA_000988465.1 | *Escherichia coli* |
| SQ37 | GCA_000988355.1 | *Escherichia coli* |
| SQ88 | GCA_000988385.1 | *Escherichia coli* |
| SRCC 1675 | GCA_001612495.1 | *Escherichia coli* |
| SS17 | GCA_000730345.1 | *Escherichia coli* |
| SS52 | GCA_000803705.1 | *Escherichia coli* |
| ST2747 | GCA_000599665.1 | *Escherichia coli* |
| ST2747 | GCA_000599685.1 | *Escherichia coli* |
| ST2747 | GCA_000599705.1 | *Escherichia coli* |
| ST540 #1 | GCA_000597845.1 | *Escherichia coli* |
| ST540 #2 | GCA_000599625.1 | *Escherichia coli* |
| ST540 #3 | GCA_000599645.1 | *Escherichia coli* |
| ST648 | GCA_001485455.1 | *Escherichia coli* |
| TW14359 | GCA_000022225.1 | *Escherichia coli* |
| uk_P46212 | GCA_001469815.1 | *Escherichia coli* |
| UM146 | GCA_000148605.1 | *Escherichia coli* |
| UMNK88 | GCA_000212715.2 | *Escherichia coli* |
| UTI89 | GCA_000013265.1 | *Escherichia coli* |
| VR50 | GCA_000968515.1 | *Escherichia coli* |
| W #1 | GCA_000184185.1 | *Escherichia coli* |
| W #2 | GCA_000258145.1 | *Escherichia coli* |
| WS4202 | GCA_001307215.1 | *Escherichia coli* |
| Xuzhou21 | GCA_000262125.1 | *Escherichia coli* |
| YD786 | GCA_001442495.1 | *Escherichia coli* |
| ZH063 | GCA_001577325.1 | *Escherichia coli* |
| ZH193 | GCA_001566675.1 | *Escherichia coli* |
| 11E12 | GCA_002260745.1 | *Klebsiella aerogenes* |
| 1019_EAER | GCA_001053595.1 | *Klebsiella aerogenes* |
| 1020_EAER | GCA_001052095.1 | *Klebsiella aerogenes* |
| 1277_EAER | GCA_001052565.1 | *Klebsiella aerogenes* |
| 1278_EAER | GCA_001054275.1 | *Klebsiella aerogenes* |
| 1282_EAER | GCA_001053235.1 | *Klebsiella aerogenes* |
| 151_EAER | GCA_001054405.1 | *Klebsiella aerogenes* |
| 225_EAER | GCA_001071835.1 | *Klebsiella aerogenes* |
| 86_EAER | GCA_001058645.1 | *Klebsiella aerogenes* |
| AR_0007 | GCA_002796425.1 | *Klebsiella aerogenes* |
| AR_0009 | GCA_002796525.1 | *Klebsiella aerogenes* |
| AR_0018 | GCA_002796405.1 | *Klebsiella aerogenes* |
| AR_0062 | GCA_002948835.2 | *Klebsiella aerogenes* |
| AR_0161 | GCA_003071285.1 | *Klebsiella aerogenes* |
| C10 | GCA_001662765.1 | *Klebsiella aerogenes* |
| CAV1320 | GCA_001021995.1 | *Klebsiella aerogenes* |
| CDC UA0804-01 | GCA_000755545.1 | *Klebsiella aerogenes* |
| D2 | GCA_001662705.1 | *Klebsiella aerogenes* |
| D3 | GCA_001662695.1 | *Klebsiella aerogenes* |
| E20 | GCA_002946615.1 | *Klebsiella aerogenes* |
| E9 | GCA_001662715.1 | *Klebsiella aerogenes* |
| EA1509E | GCA_000334515.1 | *Klebsiella aerogenes* |
| Ea77 | GCA_001649605.2 | *Klebsiella aerogenes* |
| FDAARGOS_139 | GCA_001593585.2 | *Klebsiella aerogenes* |
| FDAARGOS_152 | GCA_001559215.2 | *Klebsiella aerogenes* |
| FDAARGOS_327 | GCA_003546885.1 | *Klebsiella aerogenes* |
| FDAARGOS_363 | GCA_002591115.1 | *Klebsiella aerogenes* |
| FDAARGOS_513 | GCA_003812185.1 | *Klebsiella aerogenes* |
| FGI35 | GCA_000383335.1 | *Klebsiella aerogenes* |
| G7 | GCA_001571545.2 | *Klebsiella aerogenes* |
| kae_122664 | GCA_901484885.1 | *Klebsiella aerogenes* |
| KCTC 2190 | GCA_000215745.1 | *Klebsiella aerogenes* |
| MGH 61 | GCA_000692155.1 | *Klebsiella aerogenes* |
| MGH 62 | GCA_000692175.1 | *Klebsiella aerogenes* |
| MGH 77 | GCA_000692195.1 | *Klebsiella aerogenes* |
| MGH 78 | GCA_000692215.1 | *Klebsiella aerogenes* |
| MGH174 | GCA_002152895.1 | *Klebsiella aerogenes* |
| NCTC10006 | GCA_900635435.1 | *Klebsiella aerogenes* |
| NCTC9735 | GCA_900637945.1 | *Klebsiella aerogenes* |
| NFIX39 | GCA_900119335.1 | *Klebsiella aerogenes* |
| PX01 | GCA_002204605.1 | *Klebsiella aerogenes* |
| SCKA020061 | GCA_002852865.1 | *Klebsiella aerogenes* |
| UCI 15 | GCA_000534335.1 | *Klebsiella aerogenes* |
| UCI 16 | GCA_000534315.1 | *Klebsiella aerogenes* |
| UCI 27 | GCA_000534255.1 | *Klebsiella aerogenes* |
| UCI 28 | GCA_000534235.1 | *Klebsiella aerogenes* |
| UCI 45 | GCA_000534135.1 | *Klebsiella aerogenes* |
| UCI 46 | GCA_000534115.1 | *Klebsiella aerogenes* |
| UCI 47 | GCA_000534095.1 | *Klebsiella aerogenes* |
| UCI 48 | GCA_000534075.1 | *Klebsiella aerogenes* |
| UCI119 | GCA_002152915.1 | *Klebsiella aerogenes* |
| UCI120 | GCA_002152925.1 | *Klebsiella aerogenes* |
| UCI89 | GCA_001030125.1 | *Klebsiella aerogenes* |
| UCI90 | GCA_001030165.1 | *Klebsiella aerogenes* |
| UCI97 | GCA_001030185.1 | *Klebsiella aerogenes* |
| UMB0295 | GCA_002871375.1 | *Klebsiella aerogenes* |
