## Supplementary figures and images for "Microevolution of acquired colistin resistance in Enterobacteriaceae from ICU patients receiving selective decontamination of the digestive tract"

### Supplemental Figure 1

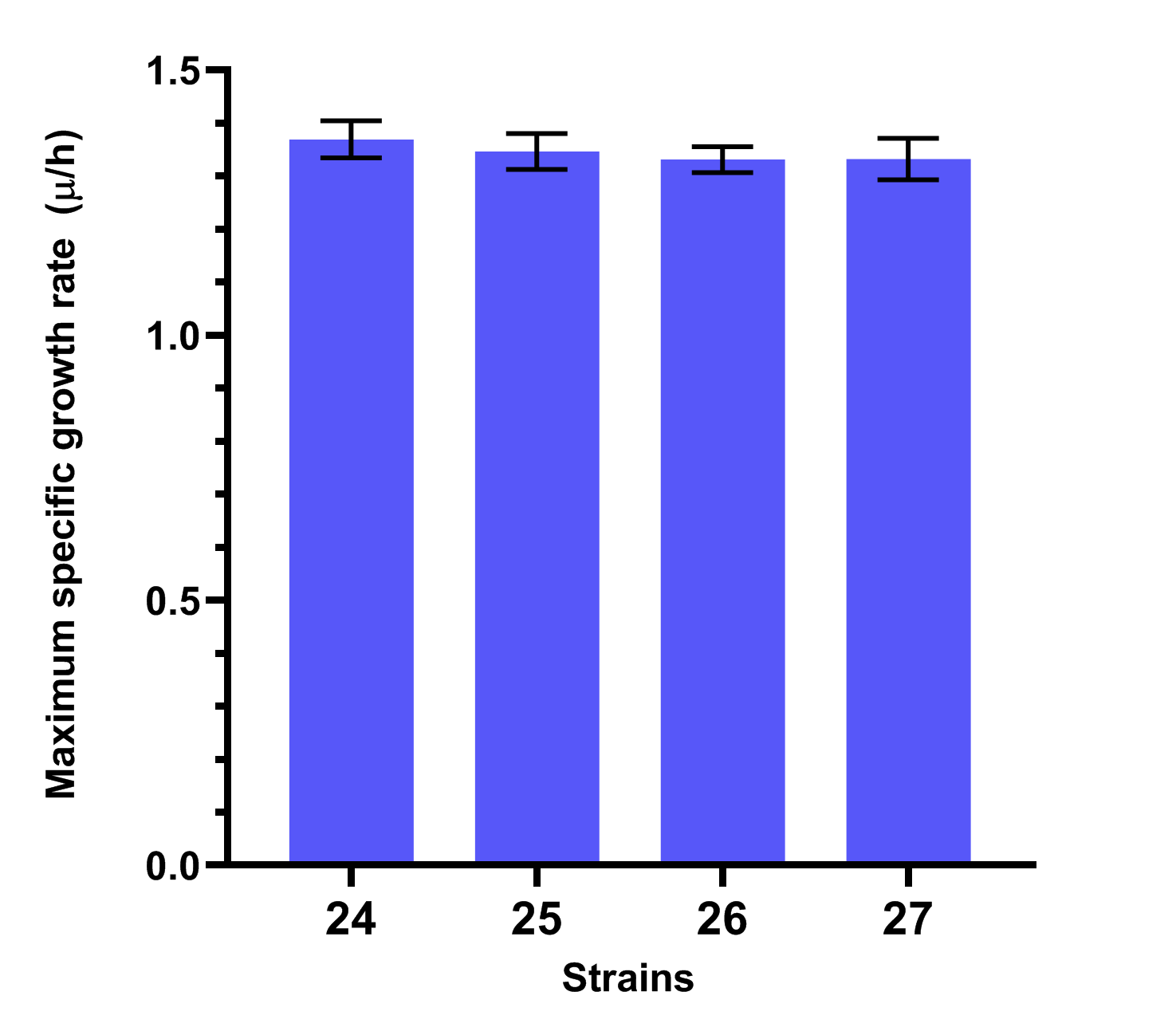
